# The number of *k*-mer matches between two *DNA* sequences as a function of *k* and applications to estimate phylogenetic distances

**DOI:** 10.1101/527515

**Authors:** Sophie Röhling, Alexander Linne, Jendrik Schellhorn, Morteza Hosseini, Thomas Dencker, Burkhard Morgenstern

## Abstract

We study the number *N*_*k*_ of length-*k* word matches between pairs of evolutionarily related DNA sequences, as a function of *k*. We show that the *Jukes-Cantor* distance between two genome sequences – i.e. the number of substitutions per site that occurred since they evolved from their last common ancestor – can be estimated from the slope of a function *F* that depends on *N*_*k*_ and that is affine-linear within a certain range of *k*. Integers *k*_min_ and *k*_max_ can be calculated depending on the length of the input sequences, such that the slope of *F* in the relevant range can be estimated from the values *F* (*k*_min_) and *F* (*k*_max_). This approach can be generalized to so-called *spaced-word* matches, where mismatches are allowed at positions specified by a user-defined binary pattern. Based on these theoretical results, we implemented a software program for alignment-free sequence comparison called *Slope-SpaM*. Test runs on real and simulated sequence data show that *Slope-SpaM* can accurately estimate phylogenetic distances for distances up to around 0.5 substitutions per position. The statistical stability of our results is improved if spaced words are used instead of contiguous words. Unlike previous alignment-free methods that are based on the number of (spaced) word matches, *Slope-SpaM* produces accurate results, even if sequences share only local homologies.

## 1 Introduction

Phylogeny reconstruction is a fundamental task in computational biology [15]. Here, a basic step is to estimate pairwise evolutionary distances between protein or nucleic-acid sequences. Under the *Jukes-Cantor* model of evolution [25], the distance between two evolutionarily related *DNA* sequences can be defined as the number of nucleotide substitutions per site that have occurred since the two sequences have evolved from their last common ancestor. Traditionally, phylogenetic distances are inferred from pairwise or multiple sequence alignments. For the huge amounts of sequence data that are now available, however, sequence alignment has become too slow. Therefore, considerable efforts have been made in recent years, to develop fast *alignment-free* approaches that can estimate phylogenetic distances without the need to calculate full alignments of the input sequences, see [20, 50, 57, 4, 26] for recent review articles. Alignment-free approaches are not only used in phylogeny reconstruction, but are also important in metagenomics [10, 39, 32] and in medical applications, for example to identify drug-resistant bacteria [5] or to classify viruses [55, 2]. In all these applications, it is crucial to rapidly estimate pairwise similarity or dissimilarity values in large sets of sequence data.

Some alignment-free approaches are based on word frequencies [40, 48] or on the length of common substrings [52, 28, 23]. Other methods use variants of the *D*_2_ distance which is defined as the number of word matches of a predefined length between two sequences [41, 53, 51, 2]; a review focusing on these methods is given in [42]. *kWIP* [37] is a further development of this concept that uses information-theoretical weighting. Most of these approaches calculate heuristic measures of sequence (dis-)similarity that are difficult to interpret. At the same time, alignment-free methods have been proposed that can accurately estimate phylogenetic distances between sequences based on stochastic models of DNA or protein evolution, using the length of common substrings [22, 35] or so-called *micro alignments* [54, 21, 30, 29, 12].

Several authors have proposed to estimate phylogenetic distances from the number of *k*-mer matches between two sequences. The tools *Cnidaria* [1] and *AAF* [14] use the *Jaccard index* between sets of *k*-mers from two – assembled or unassembled – genomes to estimate the distance between them. In *Mash* [39], the *MinHash* [6] approach is used to reduce the input sequences to small ‘sketches’ which can be used to rapidly approximate the *Jaccard index*. *Skmer* [47] is a further improvement of this approach. In a previous paper, we proposed another way to infer evolutionary distances between DNA sequences based on the number of word matches between them, and we generalized this to so-called *spaced-word* matches [36]. Here, a *spaced-word match* is a pair of words from two sequences that are identical at certain positions, specified by a pre-defined binary pattern of *match* and *don’t-care* positions.

The distance function proposed in [36] is now used by default in the program *Spaced* [27]. Theoretically, this distance measure is based on a simple model of molecular evolution without insertions or deletions. In particular, we assumed in this previous paper that the compared sequences are homologous to each other over their entire length. In practice, the derived distance values are still reasonably accurate if a limited number of insertions and deletions is allowed, and phylogenetic trees could be obtained from these distance values that are similar to trees obtained with more traditional approaches. Like other methods that are based on the number of common *k*-mers, however, our previous approach can no longer produce accurate results if sequences share only local regions of homology.

Recently, Bromberg *et al.* published an interesting new approach to alignment-free protein sequence comparison that they called *Slope Tree* [7]. They defined a distance measure using the decay of the number of *k*-mer matches between two sequences, as a function of *k*. Trees reconstructed with this distance measure were in accordance to known phylogenetic trees for various sets of prokaryotes. *Slope Tree* can also correct for horizontal gene transfer and can deal with composition variation and low complexity sequences. From a theoretical point-of-view, however, it is not clear whether the distance measure used in *Slope Tree* is an accurate estimator of evolutionary distances.

In the present paper, we study the number *N*_*k*_ of word or spaced-word matches between two DNA sequences, where *k* is the word length or the number of *match positions* of the underlying pattern for spaced words, respectively. Inspired by Bromberg’s *Slope Tree* approach, we study the decay of *N*_*k*_ as a function of *k*. More precisely, we define a function *F* (*k*) that depends on *N*_*k*_ and that can be approximated – under a simple probabilistic model of DNA evolution, and for a certain range of *k* – by an affine-linear function of *k*. The distance between two sequences – defined as the number of substitutions per site – can be estimated from the slope of *F* in this range, we therefore call our implementation *Slope-SpaM*, where *SpaM* stands for *<u>Spa</u>ced-Word <u>M</u>atches*. In an earlier implementation, we calculated *N*_*k*_ and *F* (*k*) for many values of *k*, in order to find the relevant affine-linear range of *F* and its slope therein [44]. In the present paper, we show how two values *k*_min_ and *k*_max_ can be calculated within this range such that the evolutionary distance between the compared sequences can be estimated from the values *F* (*k*_min_) and *F* (*k*_max_). Since it suffices to calculate the number *N*_*k*_ of (spaced) word matches for only two values *k* = *k*_min_ and *k* = *k*_max_, this new implementation much faster than the previous version of *Slope-SpaM*.

Using simulated DNA sequences, we show that *Slope-SpaM* can accurately estimate phylogenetic distances. In contrast to other methods that are based on the number of (spaced) word matches, *Slope-SpaM* produces still accurate results if the compared sequences contain only local homologies. In addition, we applied *Slope-SpaM* to infer phylogenetic trees based on genome sequences from the benchmarking project *AFproject* [56].

## 2 Design and Implementation

### 2.1 The number of *k*-mer matches as a function of *k*

We are using standard notation from stringology as used, for example, in [18]. For a string or sequence *S* over an alphabet 𝓐, |*S*| denotes the length of *S*, and *S*(*i*) is the *i*-th character of *S*, 1 ≤ *i* ≤ |*S*|. *S*[*i..j*] is the (contiguous) substring of *S* from *i* to *j*. We consider a pair of DNA sequences *S*_1_ and *S*_2_that have evolved under the *Jukes-Cantor* substitution model [25] from some unknown ancestral sequence. That is, we assume that substitution rates are equal for all nucleotides and sequence positions, and that substitution events at different positions are independent of each other. For simplicity, we first assume that there are no insertions and deletions (indels). We call a pair of positions or *k*-mers from *S*_1_ and *S*_2_, respectively, *homologous* if they go back to the same position or *k*-mer in the ancestral sequence. In our model, we have a nucleotide match probability *p* for homologous positions and a background match probability *q* for non-homologous nucleotides; the probability of two homologous *k*-mers to match exactly is therefore *p*^*k*^.

In our indel-free model, *S*_1_ and *S*_2_ must have the same length *L* = |*S*_1_| = |*S*_2_|, and positions *i*_1_ and *i*_2_ in *S*_1_ and *S*_2_, respectively, are homologous if and only if *i*_1_ = *i*_2_. Note that, under this model, a pair of *k*-mers from *S*_1_ and *S*_2_is either homologous – if the *k*-mers start at the same respective positions in the sequences – or completely non-homologous, in the sense that *none* of the corresponding pairs of positions is homologous. For a pair of non-homologous *k*-mers from *S*_1_ and *S*_2_, the probability of an exact match is *q*^*k*^.

Let the random variable *X*_*k*_ be defined as the number of *k*-mer matches between *S*_1_ and *S*_2_. More precisely, *X*_*k*_ is defined as the number of pairs (*i*_1_*, i*_2_) for which

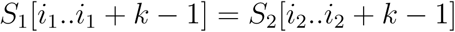

holds. There are (*L − k* + 1) possible homologous and (*L − k* + 1) · (*L − k*) possible background *k*-mer matches. The expected total number of *k*-mer matches is therefore

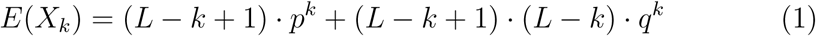

In [36], we used this equation to estimate the match probability *p* for two observed sequences using a *moment-based* approach, by replacing the expected number *E*(*X*_*k*_) by the empirical number *N*_*k*_ of word matches or spaced-word matches, respectively. Although in equation (1), an indel-free model is assumed, we could show in our previous paper, that this approach gives still reasonable estimates of *p* for sequences with insertions and deletions, as long as the sequences are globally related, *i.e.* as long as the insertions and deletions are small compared to the length of the sequences. It is clear, however, that this estimate will become inaccurate in the presence of large insertions and deletions, *i.e.* if sequences are only locally related.

Herein, we propose a different approach to estimate evolutionary distances from the number *N*_*k*_ of *k*-mer matches, by considering the *decay* of *N*_*k*_ if *k* increases. For simplicity, we first consider an indel-free model as above. From equation (1), we obtain

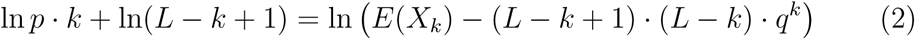

which – for a suitable range of *k* – is an approximately affine-linear function of *k* with slope ln *p*. Substituting the expected value *E*(*X*_*k*_) in the right-hand side of (2) with the corresponding *empirical* number *N*_*k*_ of *k*-mer matches for two observed sequences, we define

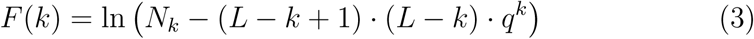

In principle, we can now estimate *p* as the exponential of the slope of *F*, as long as we restrict ourselves to a certain range of *k*. If *k* is too small, however, *N*_*k*_ will be dominated by background word matches, if *k* is too large, no word matches will be found at all. The range where we can use the slope of the function *F* to estimate the match probability *p* is discussed below.

The above considerations can be generalized to a model of *DNA* evolution with insertions and deletions and to sequences that share only *local* homologies. Let us consider two *DNA* sequences *S*_1_ and *S*_2_ of different lengths *L*_1_ and *L*_2_, respectively, that have evolved from a common ancestor under the *Jukes-Cantor* model, this time with insertions and deletions, and possibly including additional non-homologous parts of the sequences. Note that, unlike with the indel-free model with global homology, it is now possible that a *k*-mer match involves homologous as well as background nucleotide matches. Instead of deriving exact equations like (1) and (2), we therefore make some simplifications and approximations.

We can decompose *S*_1_ and *S*_2_ into indel-free pairs of ‘homologous’ sub-strings that are separated by non-homologous segments of the sequences. Let *L*_*H*_ be the – unknown – total length of the homologous substring pairs in each sequence. As above, we define the random variable *X*_*k*_ as the number of *k*-mer matches between the two sequences, and *p* and *q* are, again, the homologous and background nucleotide match probabilities. For realistic data, we also have to take into account homologies between one sequence and the *reverse complement* of the other sequence. We therefore have twice as many background *k*-mer matches than in our above simplified model. All in all, the expected number of *k*-mer matches can be roughly estimated as

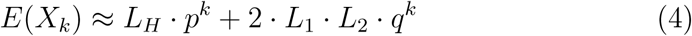

and we obtain

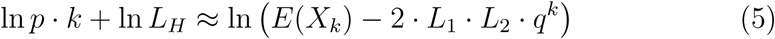

Similar as in the indel-free case, we define

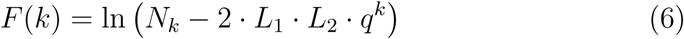

where *N*_*k*_ is, again, the empirical number of *k*-mer matches. As above, we can estimate *p* as the exponential of the slope of *F*.

An important point is that these considerations remain valid if *L*_*H*_ is small compared to *L*_1_ and *L*_2_, since *L*_*H*_ appears only in an additive constant on the left-hand side of (5). Thus, while the absolute values *F* (*k*) do depend on the extent *L*_*H*_ of the homology between *S*_1_ and *S*_2_, the *slope* of *F* only depends on *p, q, L*_1_ and *L*_2_, but not on *L*_*H*_.

### 2.2 The number of spaced-word matches

In many fields of biological sequence analysis, *k*-mers and *k*-mer matches have been replaced by *spaced words* or *spaced-word matches*, respectively. Let us consider a fixed word *P* over {0, 1} representing *match positions* (‘1’) and *don’t-care* positions (‘0’). We call such a word a *pattern*; the number of *match positions* in *P* is called its *weight*. In most applications, the first and the last symbol of a pattern *P* are assumed to be match positions. A *spaced word* with respect to *P* is defined as a string *W* over the alphabet {*A, C, G, T, \**}, of the same length as *P* with *W* (*i*) = if and only if *P* (*i*) = 0, i.e. if and only if *i* is a *don’t-care position* of *P*. Here, ‘*’ is interpreted as a *wildcard* symbol. We say that a spaced word *W* w.r.t *P* occurs in a sequence *S* at some position *i* if *W* (*m*) = *S*(*i* + *m* − 1) for all match positions *m* of *P*.

We say that there is a *spaced-word match* between sequences *S*_1_ and *S*_2_at (*i, j*) if the same spaced word occurs at position *i* in *S*_1_ and at position *j* in *S*_2_, see Fig. 1 for an example. A spaced-word match can therefore be seen as a gap-free alignment of length *P*| with matching characters at the *match positions* and possible mismatches a the |*don’t-care positions*. *Spaced-word matches* or *spaced seeds* have been introduced in database searching as an alternative to exact *k*-mer matches [8, 34, 31]. The main advantage of spaced words compared to contiguous *k*-mers is the fact that results based on spaced words are statistically more stable than results based on *k*-mers [24, 13, 10, 9, 36, 38].

**Figure 1:**
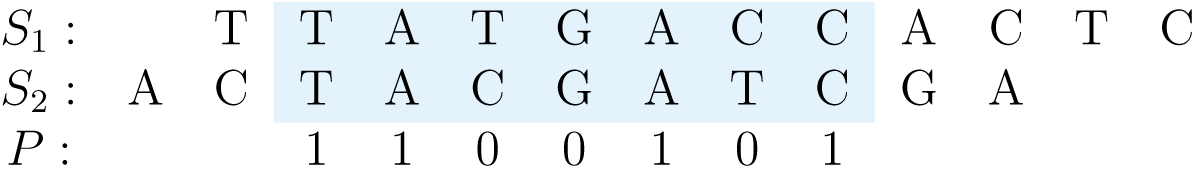
Spaced-word match between two DNA sequences *S*_1_ and *S*_2_ at (2,3) with respect to a pattern *P* = 1100101 representing *match positions* (‘1’) and *don’t-care positions* (‘0’). The same spaced word *TA* ∗ ∗*A* ∗ *C* occurs at position 2 in *S*_1_ and at position 3 in *S*_2_.

**Figure 2:**
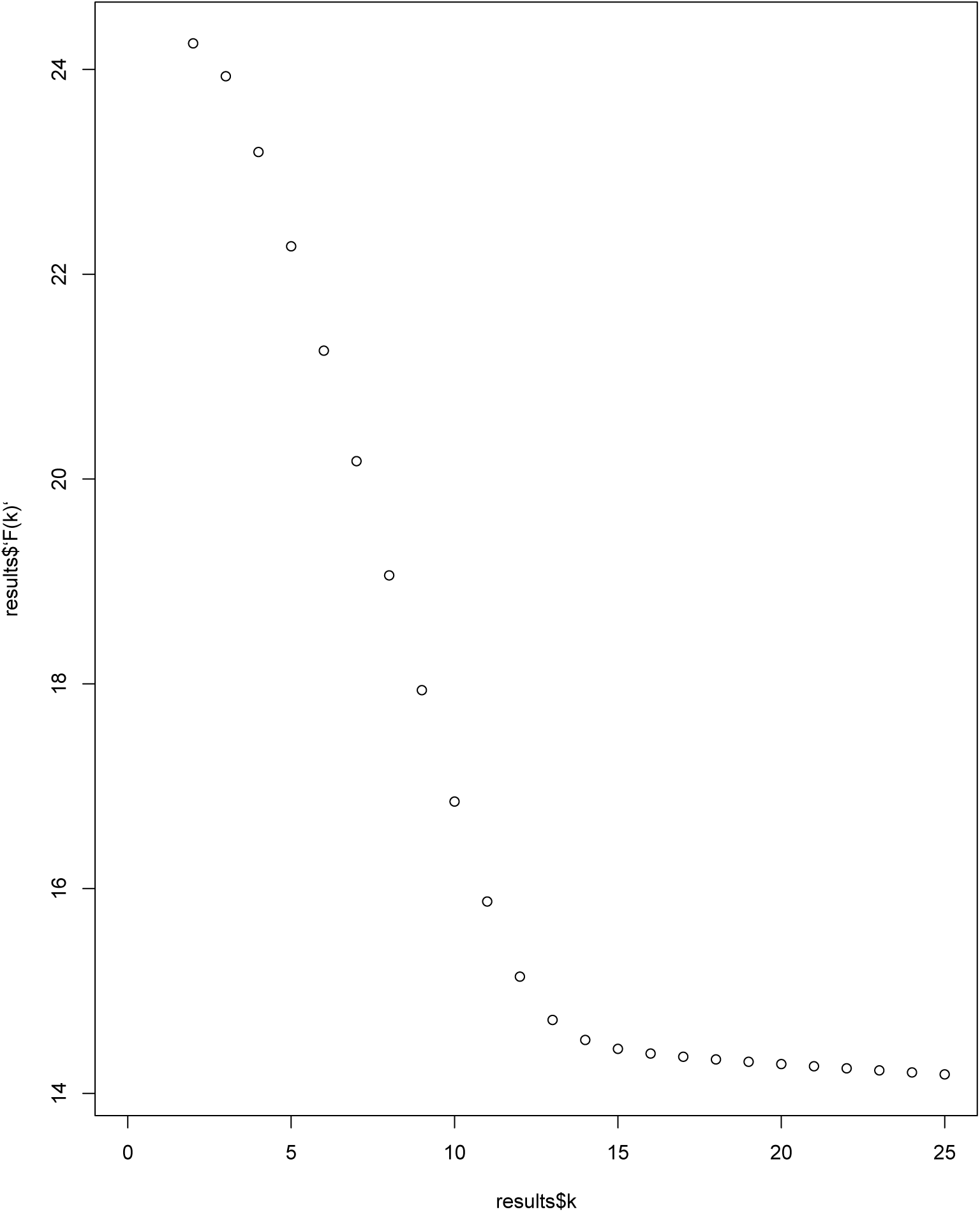
Test run on *Shigella dysenteriae 1 197* (4.44 *Mb*) and *E. coli* strain *UTI89* (5.15 *Mb*). *F* (*k*), as defined in (3), is plotted against the word length *k* for contiguous words. From the length of the sequences, the values of *k*_min_ and *k*_min_ are calculated with (14) and (15) *k*_min_ = 18 *k*_min_ = 24.

Quite obviously, approximations (4) and (5) remain valid if we define *X*_*k*_ to be the number of spaced-word matches for a given pattern *P* of weight *k*, and we can generalize the definition of *F* (*k*) accordingly: if we consider a maximum pattern weight *K* and a given set of patterns {*P*_*k*_, 1 ≤ *k* ≤ *K*} where *k* is the weight of pattern *P*_*k*_, then we can define *N*_*k*_ as the empirical number of spaced-word matches with respect to pattern *P*_*k*_ between two observed DNA sequences. *F* (*k*) can then be defined exactly as in (6), and we can estimate the nucleotide match probability *p* as the exponential of the slope of *F*.

### 2.3 The relevant affine-linear range of *F*

For simplicity, let us first assume that the two compared sequences have the same length *L*, and that they are homologous over their entire length. If, for a factor *α*, we want there to be at least *α* times more homologous than background word matches, we have to require *L⋅p*^*k*^ ≥ *α* ⋅ 2 ⋅ *L*^2^ *q*^*k*^. We therefore obtain a lower bound for *k* as

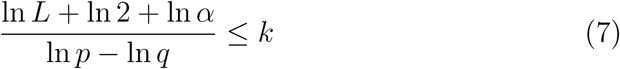

If, on the other hand, we want to have at least *β* expected word matches, we have to require *L* ⋅ *p^k^* ≥ *β*, so *k* would be upper-bounded by

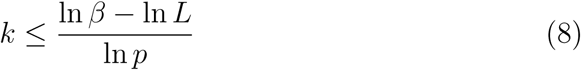

Therefore, in order to ensure that the lower bound given by (7) is smaller than the upper bound in (8), we must requi

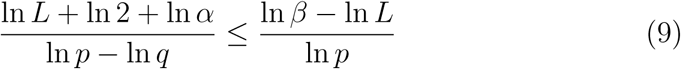

It follows that the match probability *p* must be large enough for our approach to work. More specifically, a simple transformation of (9) shows that we have to require

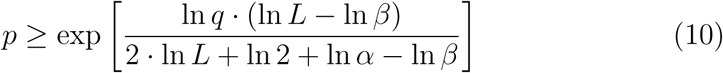

As an example, with a sequence length of 1*Gb*, a background match probability *q* = 1*/*4 and parameters *α* = *β* = 10, we would need *p >* 0.57, corresponding to a *Jukes-Cantor* distance of around 0.64 substitutions per position in order to satisfy (9) and (10), respectively.

The inequalities derived in this subsection can be easily generalized to the situation where sequences share only a local region of homology of unknown length *L*_*H*_ – as long as we can assume that the ratio between *L*_*H*_ and the total sequence length is lower-bounded by some known constant.

### 2.4 Implementation

It is well known that the set of all *k*-mer matches between two sequences can be calculated efficiently by lexicographically *sorting* the list of all *k*-mers from both sequences, such that identical *k*-mers appear as contiguous *blocks* in this list. This standard procedure can be directly generalized to spaced-word matches.

To estimate the slope of *F*, one has to take into account that the values *F* (*k*) can be used only within a certain range of *k*, as explained above. If *k* is too small, *N*_*k*_ and *F* (*k*) are dominated by the number of background (spaced) word matches. If *k* is too large, on the other hand, the number of homologous (spaced) word matches becomes small, and for *N_k_ <* 2 ⋅ *L*_1_⋅*L*_2_⋅*q*^*k*^, the value *F* (*k*) is not even defined. For two input sequences, we therefore need to identify a range *k*_min_*,…, k*_max_ in which *F* is approximately affine linear. To find such a range, we want to use inequalities (7) and (8).

There seems to be a certain difficulty in this approach, since the the inequalities that we want to use to estimate the match probability *p* depend not only on the length of the sequences and the background match probability *q*, but also on the probability *p* that we want to estimate. It is easy to see, however, that the left-hand side of (7) is monotonically *decreasing* as a function of *p*, while the right-hand side of (8) is monotonically *increasing*. We can therefore calculate *k*_min_ and *k*_max_ for a fixed small probability *p*′ and obtain values *k*_min_ and *k*_max_ that work for any match probability *p > p*′, since these values are guaranteed to be within the affine-linear region that we want to use to estimate *p*. For a suitable small match probability *p*′, we therefore define

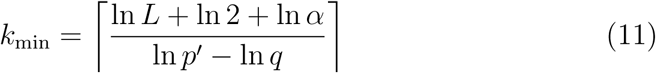

and

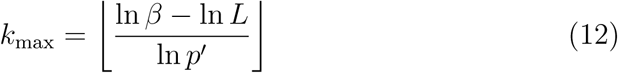

and estimate the slope of *F* as

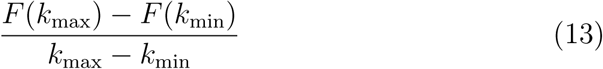

The match probability *p* can then be estimated as the exponential of this slope, provided that *p ≥ p* ′ holds.

In our test runs, we obtained good results with parameter values *α* = *β* = 1 and with two different values for *p*′, namely of *p*′ = 0.6 in (11) and *p*′ = 0.53 in (12), so are using these values by default. With a background match probability of *q* = 0.25, we therefore obtain default values of

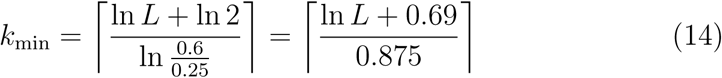

and

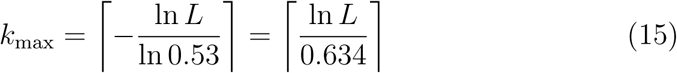

where *L* is the *average* length of the compared sequences. With these values, we then estimate *p* as

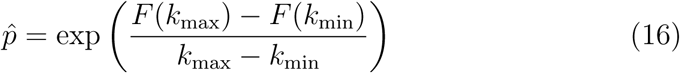

with the function *F* defined as in (3). Finally, we apply the usual *Jukes-Cantor* correction to the estimated match probability 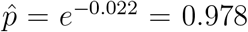, in order to estimate the *Jukes-Cantor* distance between the sequences, e.g. the number of substitutions per position that have occurred since they diverged from their last common ancestor.

## 3 Results

As a first example, to demonstrate how our approach works, we compared the genomes of *Shigella dysenteriae 1 197* (4.44 *Mb*) and *E. coli* strain *UTI89* (5.15 *Mb*). In Fig.2, *F* (*k*) is plotted against the word length *k* for contiguous words. By inserting the average sequence length *L* into (14) and (15), we obtain *k*_min_ = 19 and *k*_max_ = 24. With these values, we can calculate the slope of *F* in the relevant range as

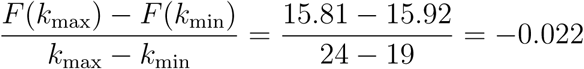

and we obtain an estimated match probability of 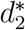. This corresponds to an average number of 0.022 substitution per position according to the *Jukes-Cantor* correction. With *Filtered Spaced Word Matches (FSWM)*, as a comparison, we obtained a distance of 0.023 substitutions per position for this pair of genomes.

### 3.1 Distances between simulated DNA sequences

For a more systematic evaluation, we generated simulated pairs of *DNA* sequences with *Jukes-Cantor* distances *d* between 0.05 and 1.0 substitutions per sequence position. For each value of *d*, we first generated 20,000 sequence pairs of length *L* = 100, 000 each, and we estimated their distances with *Slope-SpaM*. Here, we ran the program (*a*) with *k*-mers and (*b*) with spaced words based on randomly generated patterns with a probability of 0.5 for match positions. In Fig. 3, the average estimated distances are plotted against the real distances; standard deviations are represented as error bars. Our estimates are fairly accurate for distances up to around 0.5 substitutions per site, but become statistically less stable for larger distances. We did the same test runs with simulated sequences of length *L* = 1, 000, 000; the results of these runs are shown in Fig. 4. In both cases, the results were more accurate and statistically more stable when spaced words were used instead of contiguous *k*-mers.

**Figure 3:**
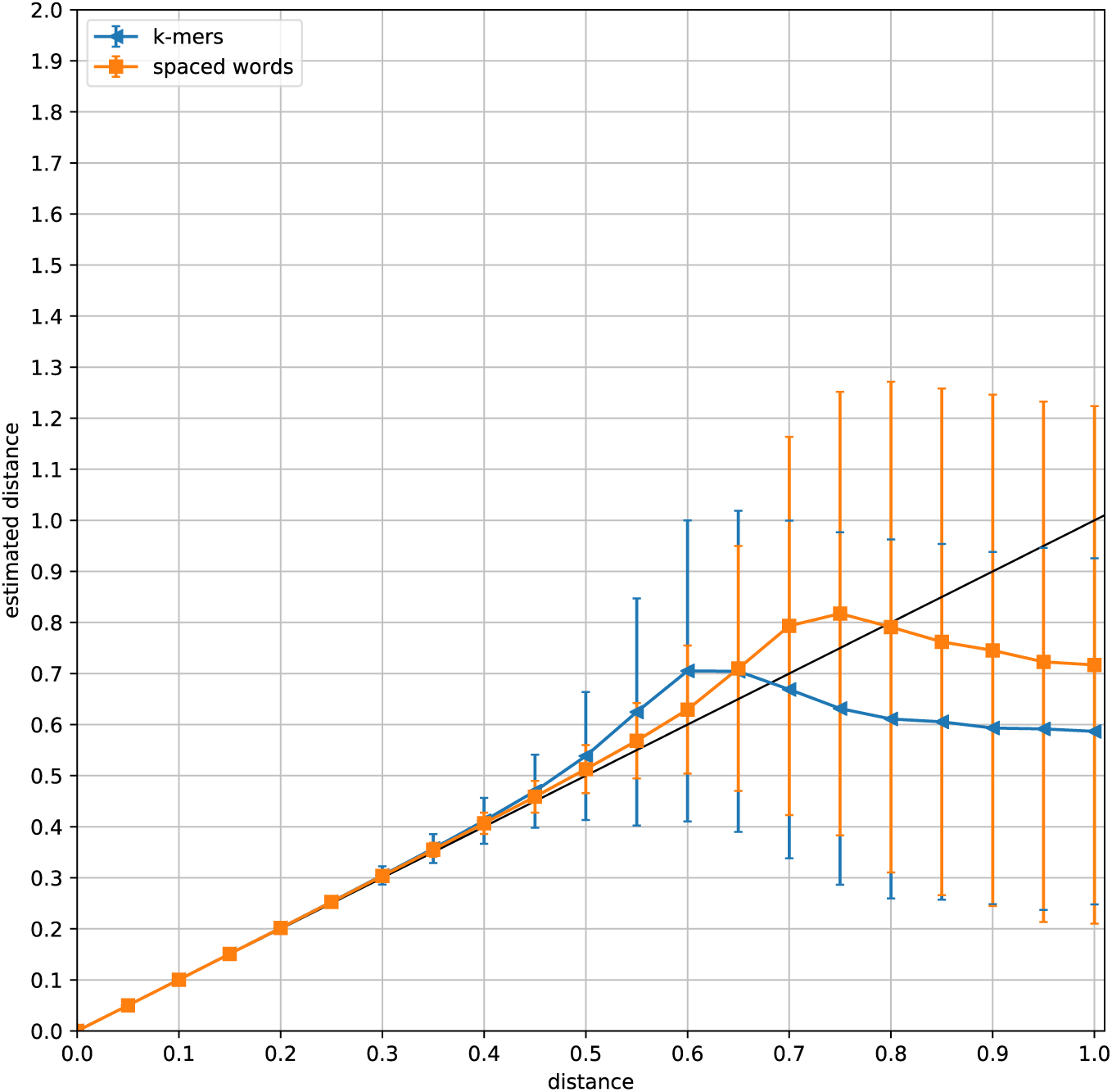
For distance values *d* between 0.05 and 1.0, we generated pairs of simulated *DNA* sequences of length *L* = 100 *kb* with a *Jukes-Cantor* distance *d*, i.e. with an average of *d* substitutions per sequence position. Distances between the sequences were estimated with *Slope-SpaM*, (*a*) based on *k*-mers and (*b*) based on spaced words with random patterns with a probability of 0.5 for a *match position* at each position. For each value of *d*, 20,000 sequence pairs were generated, their average estimated distances are plotted against the real distances. Standard deviations are shown as error bars.

**Figure 4:**
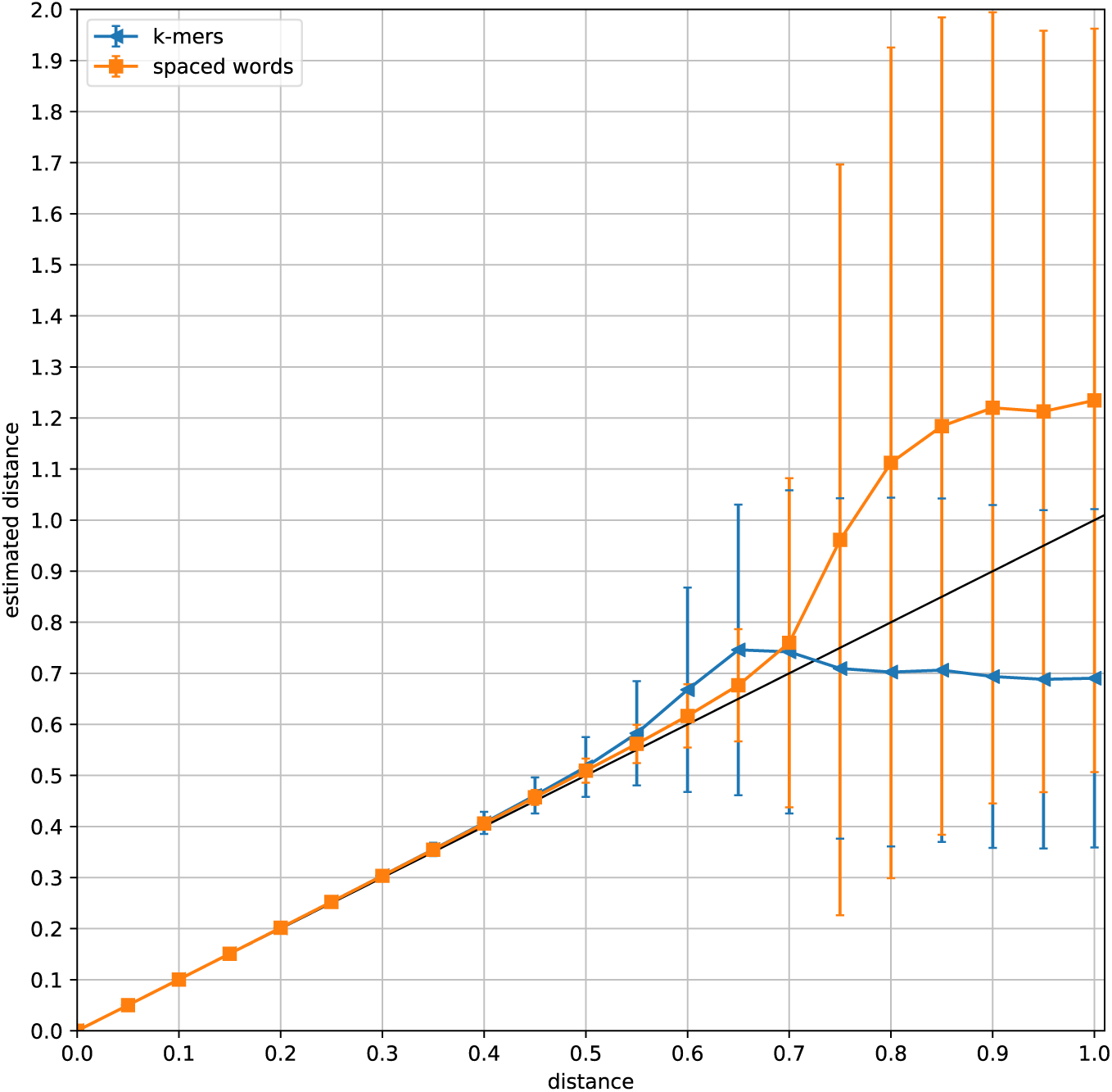
Estimated vs. real distances for pairs of simulated sequences as in Fig. 3, but with sequences of length *L* = 1*Mb*.

### 3.2 Distances between sequences with local homologies

As shown theoretically in the section *Design and Implementation*, our distance estimator is still applicable if the compared sequences share only local regions of homology, in contrast to other approaches that estimate phylogenetic distances from the number of word or spaced-word matches. To verify this empirically, we generated pairs of semi-artificial sequences consisting of local homologies, embedded in non-related random sequences of varying length. We then estimated phylogenetic distances between these sequences with *Slope-SpaM*, as well as with *Mash, Skmer* and *Spaced*, three other alignment-free methods that are also based on the number of word or spaced-word matches. To see how these tools are affected by the presence of non-homologous sequence regions, we compared the distance values estimated from the semi-artificial sequences to the distances estimated from the original sequences by the same tool.

To generate these sequence sets, we started with a set of 19 homologous genomic sequences from different strains of the bacterium *Wolbachia* from Gerth *et al.* [17]. These sequences contain 252 genes; the length of the sequences varies between 165*kb* and 175*kb* in the different strains. We then generated 9 additional data sets by adding unrelated *i.i.d.* random sequences of varying length to the left and to the right of each of the homologous genome sequences. This way, we generated 10 sets of sequences where the proportion of homology was 1.0 in the first set – the original sequences without added random sequences –, 0.9 in the second set, 0.8 in the third set, *…*, and 0.1 in the 10-th set. Within each of these 10 sequence sets, we estimated all 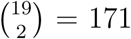 = 171 pairwise distances with the four programs that we evaluated. To find out to which extent these programs are affected by adding the non-related random sequences, we calculated for each data set and for each sequence pair the *ratio* between the estimated distance value and the distance estimated for the respective original, fully homologous sequence pair. The average ratio of these two distance values over all 171 sequence pairs is plotted against the proportion of homology in the semi-artificial sequences in Fig. 5.

**Figure 5:**
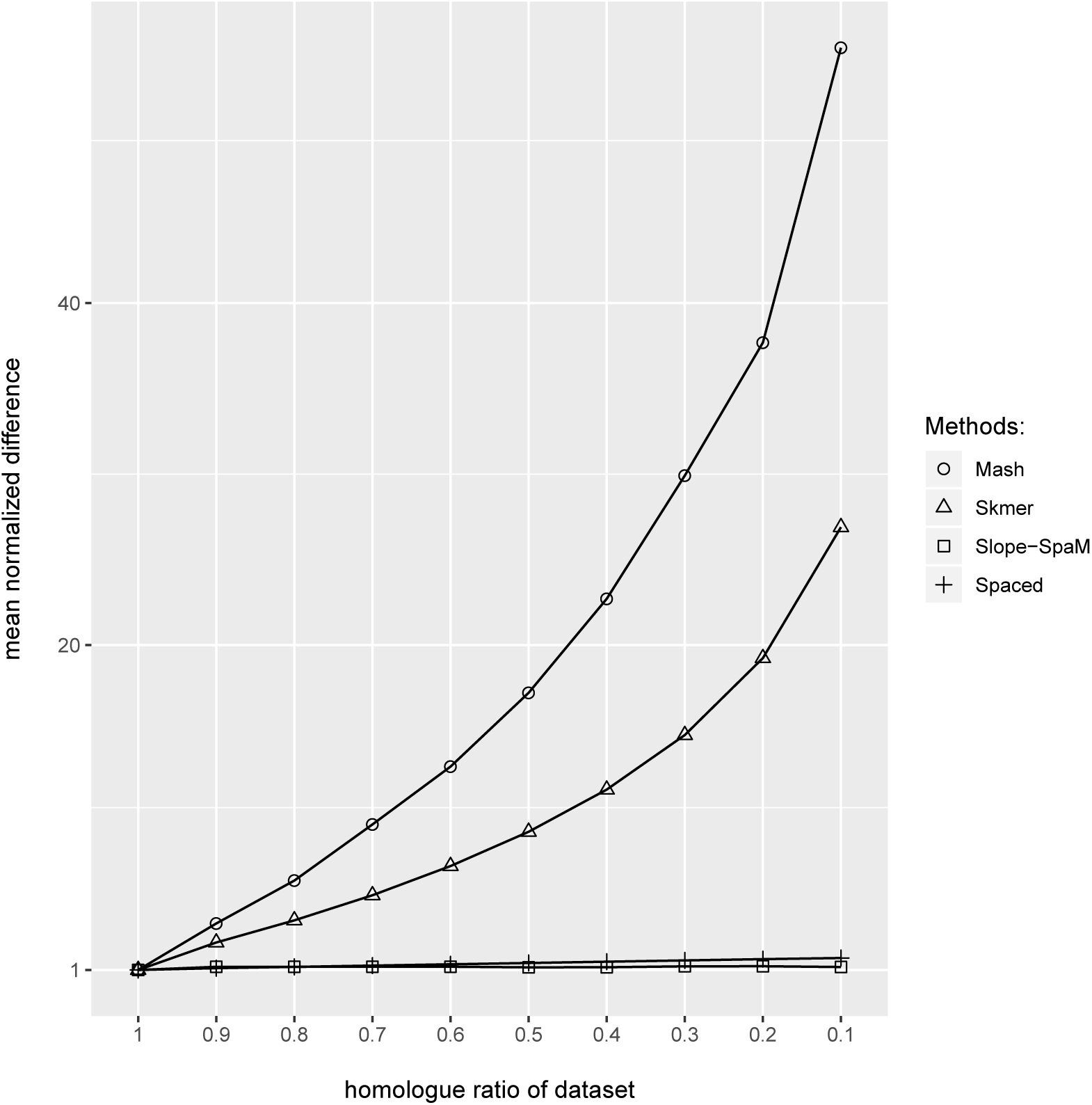
Distances estimated by four alignment-free programs between *semi-artificial* sequences consisting of 252 homologous genes from 19 strains of *Wolbachia*, embedded in non-related random sequences. Ten data sets were generated by adding random sequences of different lengths. The *x*-axis is the *proportion* of the homologous sequence within the semi-artificial sequences, the *y*-axis is the *ratio* between the distance estimated from the semi-artificial sequences and the distances estimated from the original, homologous gene sequences, see the main text for more details.

As can be seen, distances calculated with *Mash* are heavily affected by including random sequences to the original homologous sequences. For the 10-th data set, where the original homologous gene sequences account for only 10% of the generated sequences, and 90% of the sequences are unrelated random sequences, the distances calculated by *Mash* are, on average, more than 50 times as high as for the original gene sequences that are homologous over their entire length. *Skmer* and *Spaced* are also affected by the added random sequences, although to a lesser extent. By contrast, *Slope-SpaM* was hardly affected at all by the presence of the non-homologous sequences.

### 3.3 Phylogenetic tree reconstruction

In addition to the above artificial and semi-artificial sequence pairs, we used sets of real genome sequences with known reference phylogenies from the as benchmark data for phylogeny reconstruction *AFproject* [56]. *AFproject* is a collaborative project to systematically benchmark and evaluate software tools for alignment-free sequence comparison in different application scenarios. The web page of the project provides a variety benchmark sequence sets, and the output of arbitrary alignment-free methods on these sequences can be uploaded to the server. This way, the quality of these methods can be evaluated and compared to a large number of other alignment-free methods, among them the state-of-the-art methods.

To evaluate *Slope-SpaM*, we downloaded all data sets from the *AFproject* server that are relevant for our study, namely five sets of full-genome sequences that are available in the categories *genome-based phylogeny* and *horizontal gene transfer*: (1) a set of 29 *E.coli/Shigella* genomes [54], (2) a set of 14 plant genomes [19], (3) a set of 25 mitochondrial genomes from different fish species of the suborder *Labroidei* [16], (4) another set of 27 *E.coli/Shigella* [49] and (5) a set of 8 genomes of different strains of *Yersinia* [11]. These data sets have been used previously by developers of alignment-free tools to benchmark their methods.

We ran *Slope-SpaM* on the above sets of genomes and uploaded the obtained distance matrices to the *AFproject* web server for evaluation. For these categories of benchmark data, the *AFproject* server calculates phylogenetic trees from the produced distance matrices using *Neighbor Joining* [46]. It then compares the obtained trees to trusted reference trees of the respective data sets under the normalized *Robinson-Foulds (nRF)* metric. That is, the standard *Robinson-Foulds (RF)* distance [43] that measures the dissimilarity between two tree topologies is divided by the maximal possible *RF* distance of two trees with the respective number of leaves; the *nRF* distances can therefore take values between 0 and 1. On the *AFproject* server, the benchmark results are ranked in order of increasing *nRF* distance, *i.e.* in order of decreasing quality. The results of our test runs with the *AFproject* benchmark sequences are shown in Table 1.

**Table 1:** Test results on five sets of genome sequences from *AFproject*. Pairwise distance values calculated with different alignment-free methods were used as input for *Neighbor-Joining* [46]; the table contains *normalized Robinson-Foulds* distances between the resulting trees and reference trees. Thus, the smaller the values are, the more similar are the produced trees to the reference trees. The table compares the results from *Slope-SpaM* to the median results over 74 methods from the *AFproject* study. Three of the best performing programs in this study are shown (all results, except for *Slope-SpaM*, are taken from [56]).

|  | Genome-Based<br>Phylogeny |  |  | Horizontal<br>Gene Transfer |  |
| --- | --- | --- | --- | --- | --- |
|  | *mtDNA* | *E. coli* | Plants | *E. coli* | *Yersinia* |
| # genomes | 25 | 29 | 14 | 27 | 8 |
| *Slope-SpaM* (*k*-mers) | 0.32 | 0.50 | 1.00 | 0.58 | 0.20 |
| *Slope-SpaM* (spaced words) | 0.09 | 0.35 | 0.64 | 0.50 | 0.80 |
| median *AFproject* | 0.09 | 0.54 | 0.64 | 0.50 | 0.00 |
| *FFP* | 0.09 | 0.23 | 0.43 | 0.21 | 0.80 |
| *co-phylog* | 0.09 | 0.12 | 0.23 | 0.08 | 0.80 |
| *Mash* | 0.05 | 0.15 | 0.14 | 0.12 | 0.80 |
| *Skmer* | 0.09 | 0.15 | 0.25 | 0.17 | 0.80 |

It should be mentioned that the performance of the compared methods on the set of eight *Yersinia* genomes is in stark contrast to their performance on the other four data sets. This has also been observed for other methods evaluated in *AFproject*: methods that performed well on other data sets, performed badly on *Yersinia* and vice versa. It may therefore be advisable to look at this data set and the reference phylogeny used in *AFproject* in more detail.

### 3.4 Program run time

Table 2 shows the run time of *Slope-SpaM* on three of the data sets from *AFproject* that we used in the previous subsection. We used the set of 25 mitochondrial genomes from different fish species (total size 412 *kb*), the set of 29 *E. coli* genomes (total size 138 *Mb*) and the set of 14 plant genomes (total size 4.58 *Gb*).

**Table 2:**
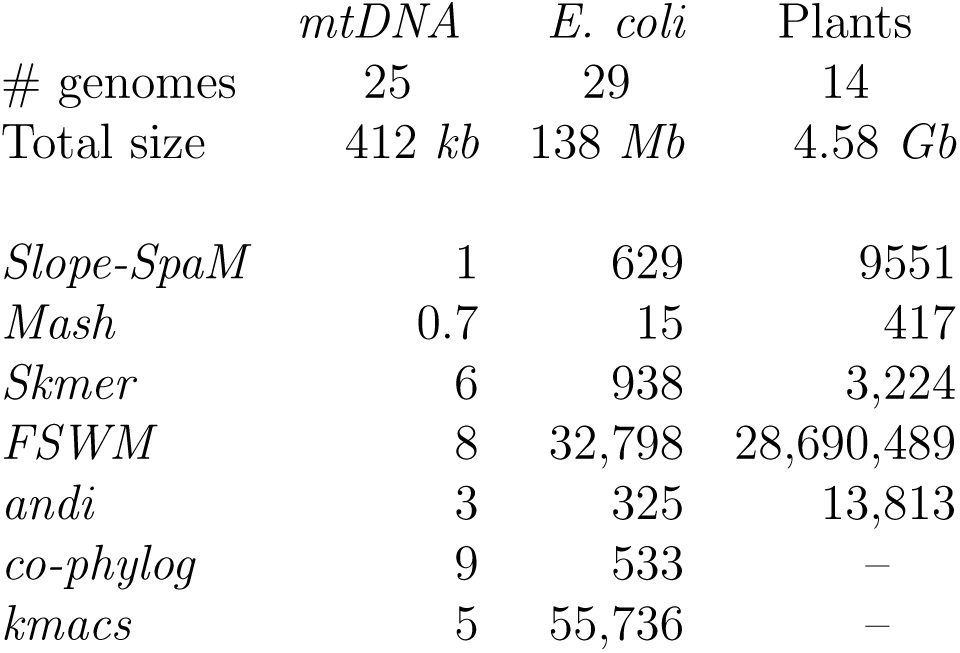
Runtime in seconds of *Slope-SpaM* (with spaced words) and six other alignment-free programs on three different sets of genomes that were used as benchmark data in *AFproject* [56]. On the largest data set, the set set of plant genomes, *co-phylog* and *kmacs* were unable to produce results.

## 4 Availability and Future Directions

A number of alignment-free methods have been proposed in recent years that estimate the nucleotide match probability *p* in an (unknown) alignment of two DNA sequences from the number *N*_*k*_ of word or *k*-mer matches for a fixed word length *k*. This can be done by comparing *N*_*k*_ to the total number of words in the compared sequences [39] or – equivalently – to the length of the sequences [35]. A certain draw-back of these approaches is that they assume that the compared sequences are homologous to each other over their entire length. Unfortunately, these methods cannot be easily applied to sequences that share only local homologies with each other. If two sequences share only a limited region of homology and are unrelated outside this region, a given number *N*_*k*_ of word matches could indicate a high match probability *p* in an alignment of those homologous regions – while the same number of word matches would correspond to a lower *p* if the homology would extend over the entire length of the sequences.

In the present paper, we introduced *Slope-SpaM*, an alternative approach to estimate the match probability *p* between two DNA sequences – and thereby their *Jukes-Cantor* distance – from the number of word matches.

The main difference between *Slope-SpaM* and previous methods is that, instead of using only one single word length *k*, our program considers a function *F* such that the values *F* (*k*) depend on the number *N*_*k*_ of word matches of length *k*. We showed in that there is a certain range of *k* where *F* is affine-linear, and the nucleotide match probability *p* in an alignment of the input sequences can be estimated from the slope of *F*. Further, we showed that one can calculate two values *k*_min_ and *k*_max_ within the relevant affine-linear range, such that it suffices to calculated *F* (*k*_min_) and *F* (*k*_max_) in order to calculate the slope of *F* in this range. The runtime of our program is therefore essentially determined by the time required to calculate the number of *k*-mer matches for only two values of *k*. From the definition of *F*, it is easy to see that the slope of *F*, and therefore the estimated match probability *p*, are not affected if the compared sequences share only local homologies. This was confirmed in our test runs with semi-artificial sequences, obtained by adding non-related random sequences to truly homologous sequences. We generalized this approach to *spaced-word matches*, i.e. word matches with possible mismatches at pre-defined positions, specified by a binary pattern of *match* and *don’t-care* positions.

To evaluate phylogenetic distances estimated by our approach, we used simulated DNA sequences with known *Jukes-Cantor* distances. For these data, we could show that distance estimates obtained with *Slope-SpaM* are highly accurate for distances up to around 0.5 substitutions per sequence position. By using spaced words instead of contiguous *k*-mers, we could improve the statistical stability of our results. To evaluate *Slope-SpaM* and other alignment-free methods on phylogenetic tree reconstruction, we used all relevant data sets from the benchmark project *AFproject*. As in the original *AFproject* study [56], we applied *Neighbor Joining* to the distance matrices produced by the approaches that we evaluated, and we compared the resulting phylogenetic tree topologies to the reference topologies that are available from the *AFproject* web page. This is a common way of evaluating alignment-free methods. As shown in Table 1, the performance of *Slope-SpaM* on these data was in the medium range, compared to the other methods that have been evaluated in *AFproject*, if our program was used with *spaced words*. We obtained better tree topologies with *spaced words* than with contiguous words. In terms of topological accuracy of the produced trees, however, *Slope-SpaM* was outperformed by some of the top methods evaluated in the *AFproject* study.

A major advantage of *Slope-SpaM*, compared to other alignment-free methods, is its high speed. It may be possible to further improve the run time of the program by using *sketching* techniques [45], to reduce the number of (spaced-)word matches that are to be considered. Conversly, if the number of word matches is too small for short or distantly related sequences, spaced-word matches based on *multiple patterns* may help to increase the range of sequences where our method can be applied.

In the last few years, a number of alignment-free approaches have been proposed that are able to use unassembled short sequencing reads as input [54, 14, 39, 33, 3, 47], as a basis for sequence clustering [1], for phylogenetic tree reconstruction [**?**] or for phylogenetic placement [3]. We are planning to evaluate systematically if *Slope-SpaM* can be used to find the position of unassembled reads in existing phylogenies, or to estimate phylogenetic distances between species based on sequencing reads instead of assembled genomes.

Since the approach implemented in *Slope-SpaM* is novel and rather different from existing alignment-free approaches, improvements may be possible by systematically analyzing the influence of the parameters of the program. In particular, there may be more sophisticated ways of finding suitable values of *k*_min_ and *k*_max_ to further improve the accuracy of *Slope-SpaM*. Also, in our study we used the *Jukes-Cantor* model, the simplest possible model for nucleotide substitutions. Using more sophisticated substitution models may also lead to improvements in future versions of the program.

The source code of our software is freely available through *GitHub*: https://github.com/burkhard-morgenstern/Slope-SpaM

## 4.1 Acknowledgements

We thank Fengzhu Sun for his useful comments on an earlier version of this manuscript, Andrzej Zielezinski and Wojchiech Karlowski for making the *AFproject* server available, Christoph Bleidorn and Micha Gerth for discussions about the *Wolbachia* genomes and Chris-André Leimeister, Christoph Elfmann and Peter Meinicke for discussions on the project.

